# HIPPOCAMPAL DNA METHYLATION, DNAm AGE AND SPATIAL MEMORY PERFORMANCE IN YOUNG AND OLD RATS

**DOI:** 10.1101/2021.05.07.443204

**Authors:** Priscila Chiavellini, Marianne Lehmann, Martina Canatelli Mallat, Joseph A. Zoller, Claudia B. Herenu, Gustavo R. Morel, Steve Horvath, Rodolfo G. Goya

**Author notes:** **Send correspondence to**: INIBIOLP, Faculty of Medicine, UNLP, CC 455, 1900 La Plata, Argentina, / 425-8988. These two authors contributed equally to this study (Priscila Chiavellini, Marianne Lehmann).

## Abstract

In humans and rats, aging is associated with a progressive deterioration of spatial learning and memory. These functional alterations are correlated with morphological and molecular changes in the brain, particularly in the hippocampus. Here, we assessed the age-related changes in the DNA methylation (DNAm) landscape in the rat hippocampus and assessed the correlation of spatial memory performance with hippocampal DNAm age in young (2.6 mo.) and old (26.6 mo.) rats.

Spatial memory performance was assessed with a modified version of the Barnes maze test. In order to evaluate learning ability as well as spatial memory retention, we assessed the time spent (permanence) by animals in goal sector 1 (GS_1_) and 3 (GS_3_) when the escape box was removed. The rat pan-tissue clock was applied to DNA methylation profiles of hippocampal tissue. The bisulfite converted genomic DNA was analyzed by Illumina Infinium HorvathMammalMethylChip40. The Horvath Mammal Methyl Chip40 assay provides quantitative measurements of DNA methylation for 22528 CpG dinucleotides that map to the Rattus norvegicus UCSC 6.0 genome. An enrichment pathway analysis revealed that neuron fate commitment, brain development, and central nervous system development were processes whose underlying genes were enriched in positively methylated CpGs. In the old rat hippocampi, the methylation levels of CpGs proximal to transcription factors associated with genes Pax5, Lbx1, Nr2f2, Hnf1b, Zic1, Zic4, Hoxd9; Hoxd10, Gli3, Gsx1 and Lmx1b, and Nipbl showed a significant regression with spatial memory performance. Regression analysis of different memory performance indices with hippocampal DNAm age was significant when data from young and old rats were taken together. The above results suggest that age-related hypermethylation of certain gene families, like Zic and Gli, may play a causal role in the decline in spatial memory in old rats. Hippocampal DNAm age seems to be a reliable index of spatial memory performance in young and old rats.

## INTRODUCTION

In humans and rats, aging is associated with a progressive deterioration of spatial learning and memory. These functional alterations are correlated with morphological changes in the brain, particularly in the hippocampus, a key brain region for the formation and consolidation of spatial memory **(1–3).** At molecular level, gene expression studies in aging rodents have documented significant changes in hippocampal genes related to cholesterol synthesis, inflammation, transcription factors, neurogenesis and synaptic plasticity **(4–8).** In the hippocampus of female rats, it was reported that 210 genes are differentially expressed in senile as compared with young counterparts, most of them being downregulated. RNA-Seq data showed that various genes involved in the immune response, including TYROBP, CD11b, C3, CD18, CD4 and CD74, are overexpressed in the hippocampus of aged animals **(9).**

A highly relevant molecular variable that has not been systematically assessed in the hippocampus of young and old rats is DNA methylation (DNAm). In particular, there are, to our knowledge, no documented studies on the impact of age changes in hippocampal DNA methylation on memory performance. Early ground-breaking studies performed in rat brain showed a global loss of DNA methylation during aging **(10)** but there is scanty information on the DNAm landscape in the rat hippocampus and even less about the impact of aging on this landscape.

The demonstration that the DNA methylation profile of a specific set of cytosine-guanine sites (CpG sites) is a highly accurate biomarker of biological age gave rise to the algorithms known as epigenetic clocks **(11–16).** Since epigenetic clocks seem to reflect biological age (DNAm age), we were interested in correlating a set of variables used to assess spatial memory, with hippocampal DNAm age in young and old rats. The results of the study are reported here.

## EXPERIMENTAL PROCEDURES

### Animals

Thirteen young (average age 2.6 mo.; range ± 2 days) and 11 old (26.6 mo; range ± 5 days.) female Sprague-Dawley (SD) rats weighing (X±SEM) 165 ± 5 and 250± 9 g, respectively, were used. Animals were housed in a temperature-controlled room (22 ± 2°C) on a 12:12 h light/dark cycle. Food and water were available *ad libitum*. All experiments with animals were performed in accordance with the Animal Welfare Guidelines of NIH (INIBIOLP’s Animal Welfare Assurance No A5647-01). The ethical acceptability of the animal protocols used here have been approved by our institutional IACUC (Protocol # T09-01-2013).

### Spatial memory assessment

#### Description of the Barnes maze protocol used

The modified Barnes maze protocol used in this study was based on a previously reported procedure **(3)**. It consists of an elevated (108 cm to the floor) black acrylic circular platform, 122 cm in diameter, containing twenty holes around the periphery. The holes are of uniform diameter (10 cm) and appearance, but only one hole is connected to a black escape box (tunnel). The escape box is 38.7 cm long × 12.1 cm wide × 14.2 cm in depth and it is removable. A white squared starting chamber (an opaque, 20 cm × 30 cm long, and 15 cm high, open-ended chamber) was used to place the rats on the platform. Four proximal visual cues were placed in the room, 50 cm away from the circular platform. The escape hole was numbered as hole 0 for graphical normalized representation purposes, the remaining holes being numbered 1 to 10 clockwise, and −1 to −9 counterclockwise **(Fig. 5-Panel A)**. Hole 0 remains in a fixed position, relative to the cues in order to avoid randomization of the relative position of the escape box. During the tests the platform was rotated daily. A 90-dB white-noise generator and a white-light 500 W bulb provided the escape stimulus from the platform. We used an abbreviated protocol based on three days of acquisition trials (AT), followed by a probe trial (PT) (one day after training) to assess spatial memory retention. An AT consists of placing a rat in the starting chamber for 30 s, the chamber is then raised, and the aversive stimuli (bright light and high pitch noise) are switched on and the rat is allowed to freely explore the maze for 120 s. A probe trial is defined as a trial where the escape box has been removed, its purpose being to assess the exploration frequency of the empty escape hole and nearby holes. After the starting chamber is raised, the rat is given 120 s to explore and the number of explorations per hole is recorded.

#### Specific procedures used in the present study

On the day before the first trial (experimental day 0), rats underwent a habituation routine to let them get acquainted with the platform and the escape box. In each AT, rats were tested (120 s per trial) with the escape box in place, two times per day for 3 consecutive days (experimental day 1 to 3). On day 4, rats were submitted to a probe trial (PT) during 120 s without an escape box. In order to eliminate olfactive clues from the maze and the boxes, the surfaces were cleaned with 10% ethylic alcohol solution, after each trial. The behavioral performances were recorded using a computer-linked video camera mounted 110 cm above the platform. The video-recorded performances of the subjects were measured using the Kinovea v0.7.6 (http://www.kinovea.org) and Image Pro Plus v5.1 (Media Cybernetics Inc., Silver Spring, MD) software. The behavioral parameters assessed were as follows.

##### (a) Escape box latency

time (in s) spent by an animal since its release from the start chamber until it enters the escape box in the last AT.

##### (b) Hole exploration frequency

during the PT is represented by a bar chart typically bell-shaped around the escape hole. The chart shows 20 bars, each corresponding to a specific hole. For a given hole, the exploratory frequency is the average number of explorations of that hole by either all young (13) or old (11) rats.

##### (c) Goal sector (GS)

Is the area of the platform corresponding to a given number of holes. Thus GS1 is the area corresponding to hole 1; GS3, is the area corresponding to holes −1, 0, +1. The permanence time in GS3 is calculated taking the time spent by the rats in the area covered by the 3 holes during the PT.

##### (g) Goal-seeking activity

the sum of explorations for all holes in a PT, divided by 20 (i.e. mean explorations per hole).

### Hippocampus Dissection

Rats were sacrificed by decapitation, the brain was removed carefully, severing the optic nerves and pituitary stalk and placed on a cold plate. The hippocampus was dissected from cortex in both hemispheres using forceps. This dissection procedure was also performed on the anterior and posterior blocks, alternatively placing the brain caudal side up and rostral side up. After dissection, each hippocampus was sagittally halved. One half fixed in PBS buffered 4% formalin and the second one immediately placed in a 1.5 ml tube and momentarily immersed in liquid nitrogen, then stored at −80°C for DNA extraction.

### Hippocampal DNA extraction

Hippocampal DNA was extracted using an automated nucleic acid extraction platform called QIAcube HT (Qiagen) with a column-based extraction kit, QIAamp 96 DNA QIAcube HT Kit (Qiagen).

### Purification and methylation analysis of genomic DNA

DNAs from 24 hemi-hippocampi were purified using the DNeasy Blood and Tissue Kit (Qiagen). Only DNA samples with 260/280 ratios greater than 2.0 were processed for methylation analysis. The bisulfite conversion of genomic DNA was performed using the EZ Methylation Kit (Zymo Research, D5002), following the manufacturer’s instructions. The converted genomic DNA was analyzed by Illumina Infinium HorvathMammalMethylChip40 at the UNGC Core Facility. The Horvath Mammal Methyl Chip40 assay provides quantitative measurements of DNA methylation for 22528 CpG dinucleotides that map to the Rattus norvegicus UCSC 6.0 genome (and other mammalian sequences) **(17).**

### Statistical analysis of differentially methylated positions

Quality control (QC), pre-processing, and statistical analysis of methylation profiles were performed with “minfi” R/Bioconductor package. **(18)** Briefly, QC analysis was performed with getQC function and further pre-processed with the Noob/ssNoob method.

Data exploratory analysis was performed through an unsupervised hierarchical clustering analysis based on Euclidean distance of methylation profiles. In addition, the distribution of the methylation measurements in each sample was evaluated. To test differential methylation levels in each CpG, multiple hypothesis testing was performed through a linear model using the “limma” package **(19).** The positive false discovery rate was controlled with a q-value threshold of 0.05 **(20)**.

Gene ontology (GO) enrichment test for positively differentially methylated CpGs from the HorvathMammal40 array was conducted with GOmeth function of “missMethyl” package **(21).** Gene Ontology gene sets were evaluated, and the significant categories at FDR<0.05 reported.

In order to establish potential linear relationship between the methylation of the CpGs present in the statistically significant GO terms and the GS3 permanence time parameter of the Barnes Maze test, we performed linear regression.

### Determination of epigenetic age

In the present study, we used the rat pan-tissue epigenetic clock, which was one of six rat clocks previously described **(22).** Briefly, the rat pan-tissue epigenetic clock was developed by regressing chronological age on CpGs from multiple rat tissues that are known to map to the genome of *Rattus norvegicus*. Age was not transformed. Penalized regression models were created with the R function “glmnet” **(23)**. We investigated models produced by both, Ridge and Lasso “elastic net” type regression (alpha=0.5). The optimal penalty parameters in all cases were determined automatically by using a 10 fold internal cross-validation (cv.glmnet) on the training set. By definition, the alpha value for the elastic net regression was set to 0.5 (midpoint between Ridge and Lasso type regression) and was not optimized for model performance. We performed a crossvalidation scheme for arriving at unbiased (or at least less biased) estimates of the accuracy of the DNAm based pan-tissue age estimators. One type consisted of leaving out a single sample (LOOCV) from the regression, predicting an age for that sample, and iterating over all samples.

The final version of the pan-tissue epigenetic clock developed in the above-referenced study was intended for future studies of rat tissue samples and is the clock used here.

## RESULTS

### Age-Related DNA methylation differences in the rat hippocampus

The aim of this analysis was to detect age-related changes in CpG methylated positions of hippocampal DNA that could be involved in the genesis of neurodegenerative disorders associated with aging. The distribution of the methylation measurements in each sample was evaluated **(Suppl. Fig. 1).** Data exploratory analysis was performed through an unsupervised hierarchical clustering, where both age groups are distinguishable **(Suppl. Fig. 2).**

Assessment of each age group showed that the methylation levels of 7222 (32%) CpGs changed with age between young and old hippocampal samples (q-value<0.05). **(Fig. 1).** Of these, 1090 (15%) CpGs exhibited increased methylation (i.e., higher beta values), with age while 5252 (73%) CpGs showed decreased methylation levels. We identified the CpGs in context with genomic feature: Promoter, Exon, Intron, Intergenic, 5’UTR, or 3’ UTR. The distribution profile of the CpG features in the genome is shown in **Fig. 2.** The profile of the feature distribution in negatively and positively methylated CpGs is similar and both are similar to the distribution of the feature positions of the Rattus norvegicus CpGs present in the Illumina Infinium HorvathMammalMethylChip40. **(Fig. 2, inset).** The number of CpGs localized in CpG islands was 869 (12%).

**Figure 1.**
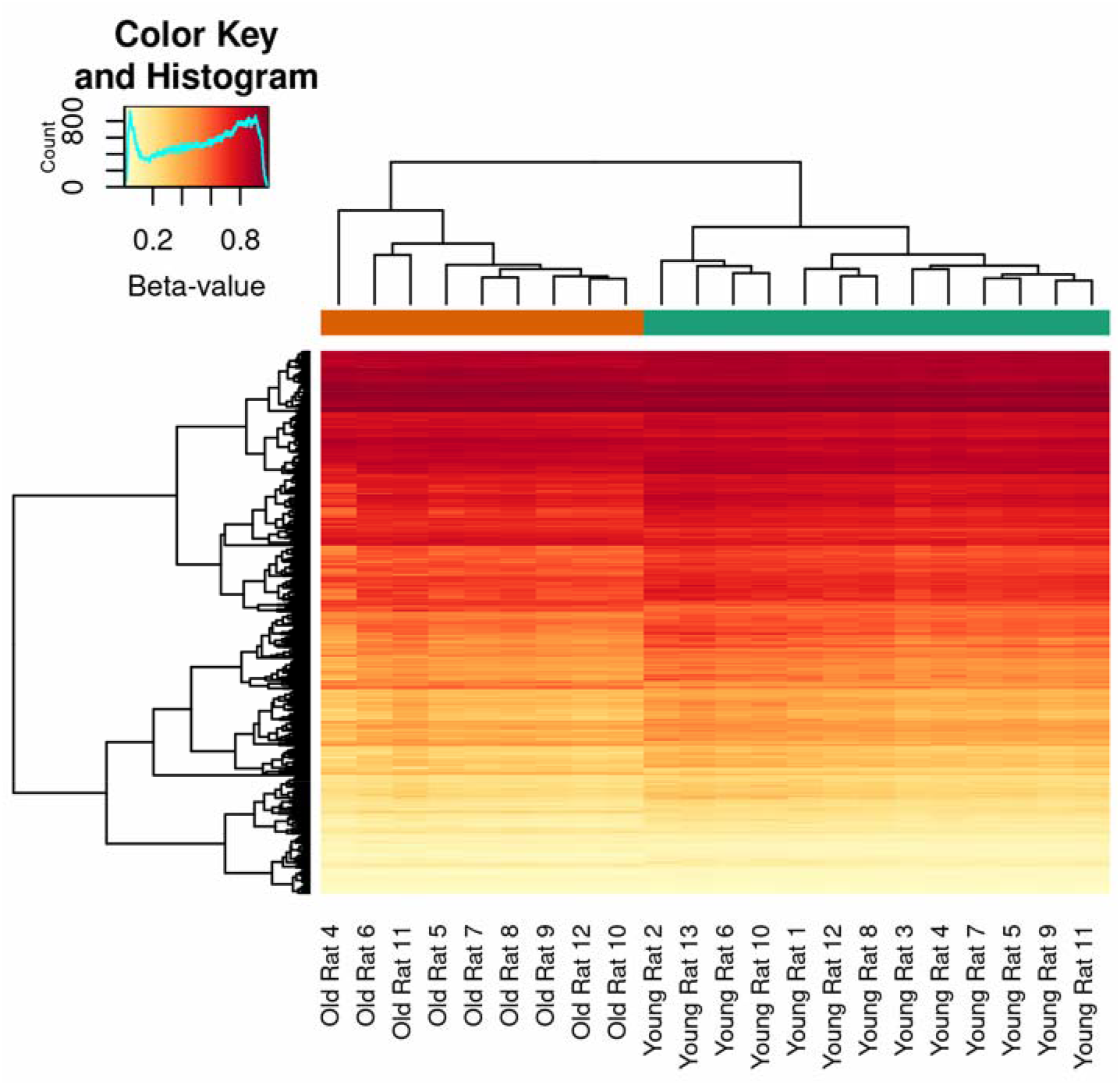
Heatmap of differentially methylated CpG sites in the old group compared with the young one. Yellow denotes CpGs with the lowest methylation levels (beta values close to 0) and red denotes CpGs with the highest methylation levels (beta values close to 1).

**Figure 2.**
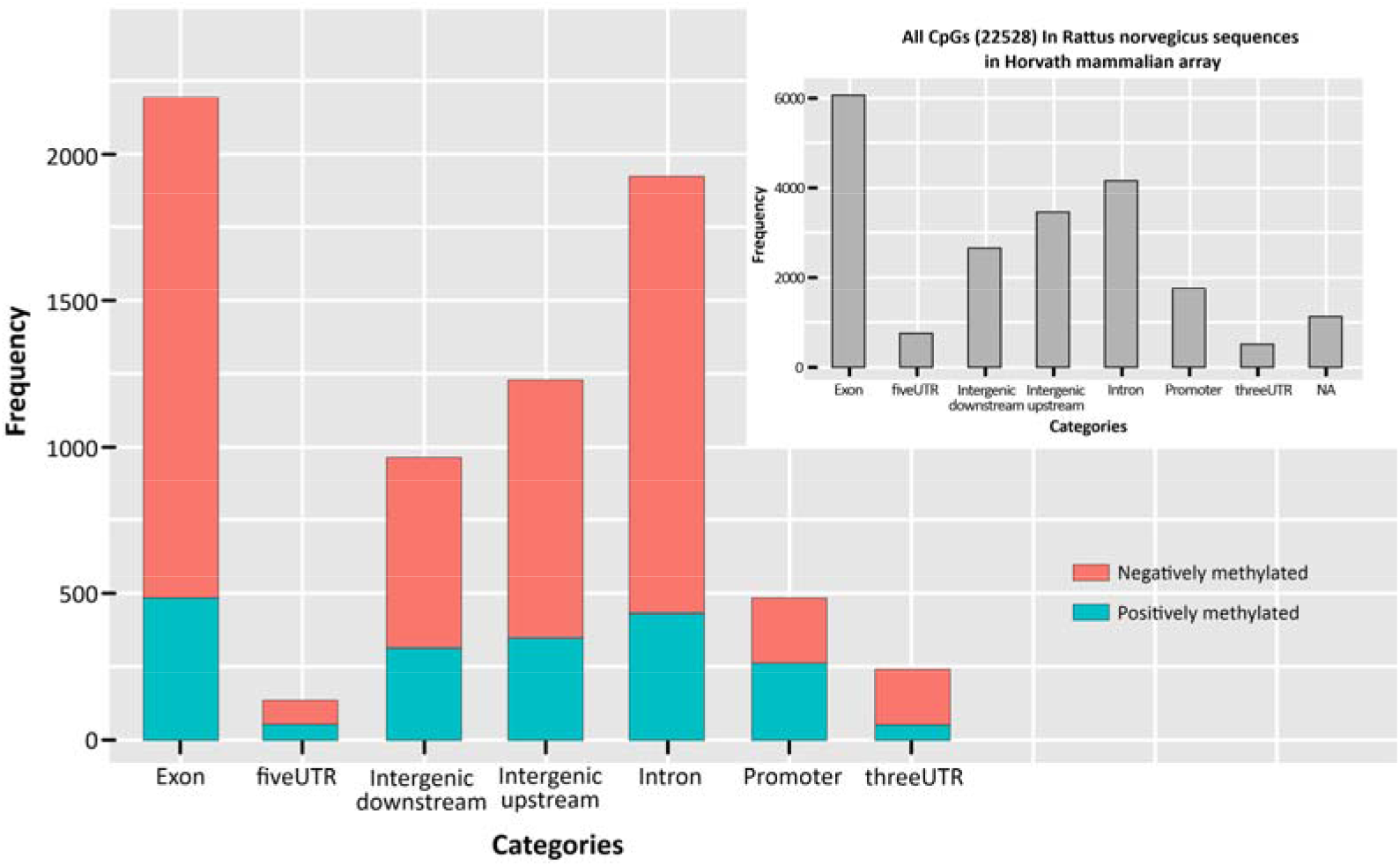
Genomic feature positions of the differentially methylated CpGs. Merged barchart of the genomic locations for the age-dependent positively methylated CpGs (in color1) and the negatively methylated CpGs (in color2). **Inset-** Feature distribution of the Rattus norvegicus sequences in the Illumina Infinium Horvath Mammal Methyl Chip40.

In order to identify potential relationships between age-related differences in DNA methylation in the nervous system (hippocampus), we performed an enrichment pathway analysis. Twenty-nine gene sets were significantly enriched in the positively methylated CpGs (FDR<0.05). Among them, three biological processes are related to the nervous system: neuron fate commitment, brain development, and central nervous system development **(Fig. 3)** On the contrary, when evaluating negatively methylated CpGs enrichment, no gene set was significantly enriched.

**Figure 3.**
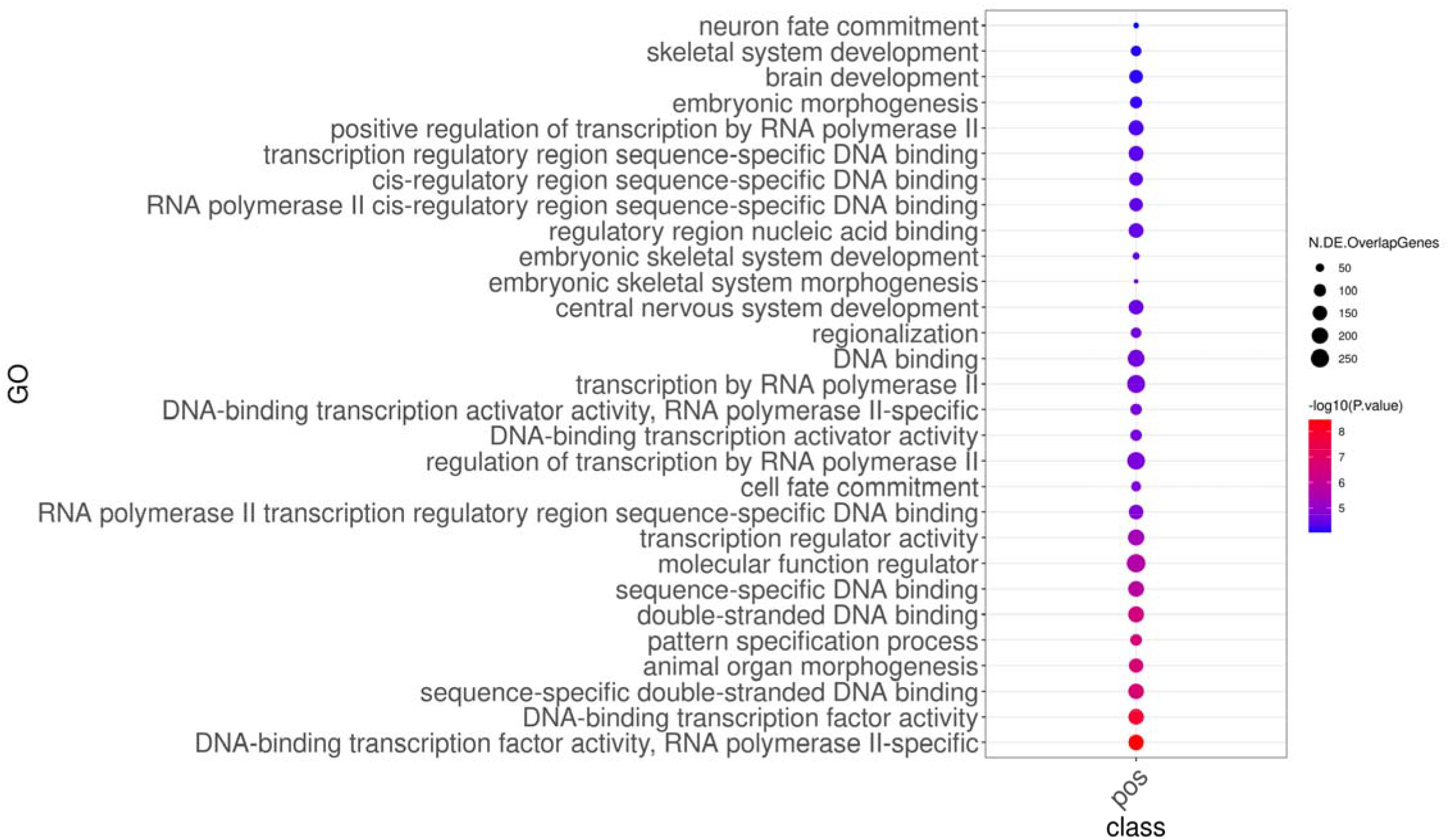
Gene ontology enrichment for positively methylated CpGs. Dot plots for significant gene sets with FDR<0.05 are displayed. Circle size represents the number of genes related to significant CpGs in a particular set.

Our last aim in the methylation landscape analysis was to identify the potential correlation between age-related changes in CpG methylation levels and hippocampus-mediated memory performance. In order to determine the relationship between age-related differences in memory performance and CpG mehylation levels near genes implicated in nervous system development, we carried out a regression analysis for such CpG sites versus GS3 permanence time index of the Barnes maze test. The reason GS3 permanence was chosen over other Barnes maze parameters is that it reliably reflects spatial memory performance as a whole (also see Discussion).

In the old rat hippocampi, we found that the methylation levels of 14 CpGs correlate negatively with the permanence time in GS3 sector **(Fig. 4).** Those CpGs are proximal to transcription factors associated with genes Pax5, Lbx1, Nr2f2, Hnf1b, Zic1, Zic4, Hoxd9; Hoxd10, Gli3, Gsx1 and Lmx1b, and Nipbl.

**Figure 4.**
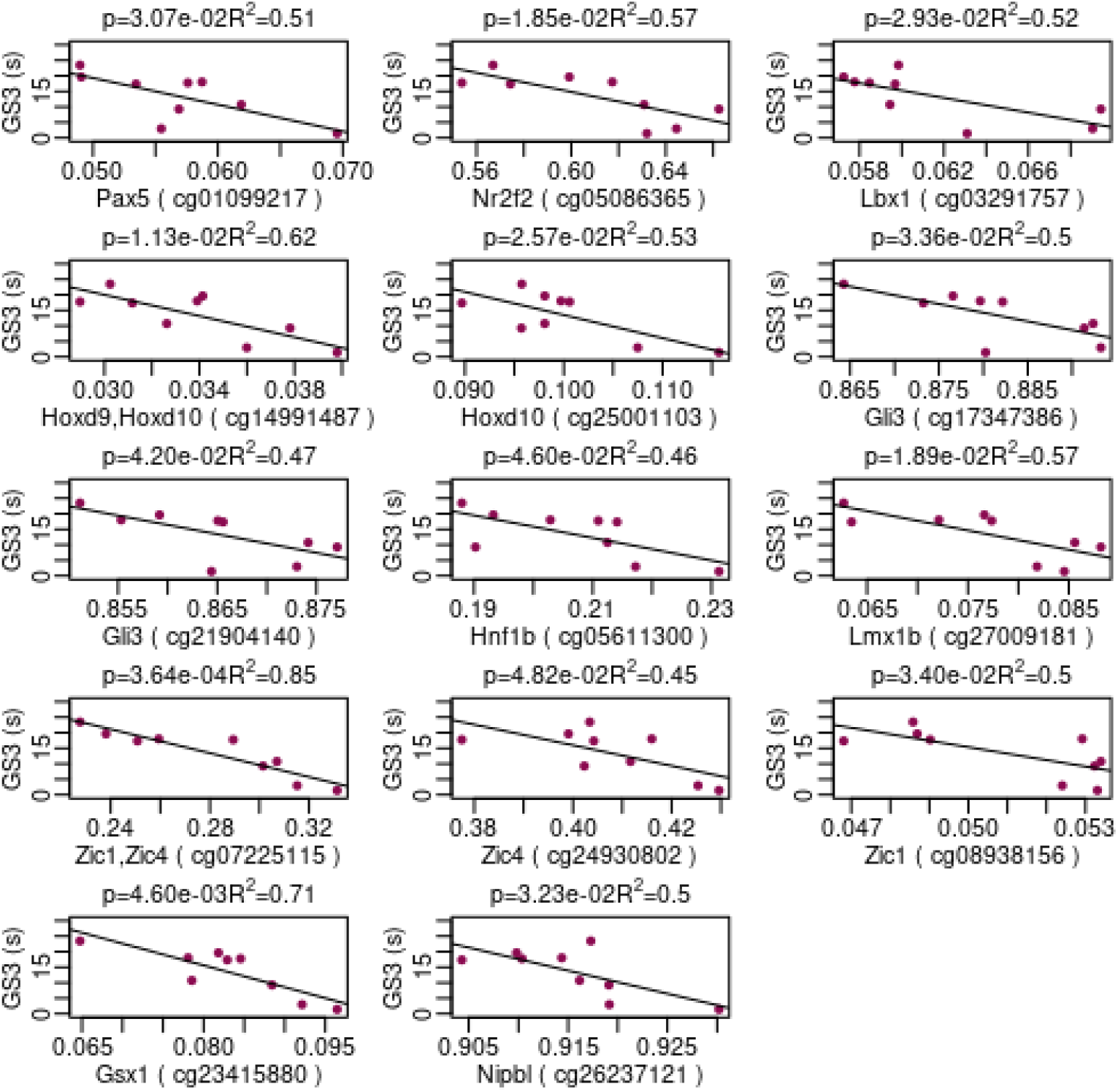
Regression plots correlating Beta-value from CpGs in the central nervous system enrichment versus permanence time in GS3 Barnes maze sector-. In old rat hippocampi, methylation levels of 14 CpGs correlate negatively with permanence time in the GS3 sector. The CpGs are proximal to transcription factors associated with genes Pax5, Lbx1, Nr2f2, Hnf1b, Zic1, Zic4, Hoxd9; Hoxd10, Gli3, Gsx1 and Lmx1b, and Nipbl.

### Spatial memory performance of young and old rats in the Barnes maze test

As expected, all spatial memory indices assessed with the Barnes maze test showed a marked decline with age. Specifically, permanence time in GS1 and GS3 dropped by 88 and 74% respectively as compared with young counterparts **(Fig . 5, panel B)**. Target-seeking activity, an index of the exploratory initiative of animals, fell by 61% as compared with young counterparts **(Fig. 5 panel C)**. During the probe trial, the hole exploratory frequency of old rats showed a flatter bell-shaped distribution around hole 0 than young counterparts. The highest difference was detected at hole 0 (93%) **(Fig. 5, panel D and inset, respectively)**. Latency to escape box, a measure of learning performance and memory retention which is inversely proportional to memory and learning performance, showed a 77% increase in the old rats versus young counterparts **(Fig. 5, panel E).**

**Figure 5.**
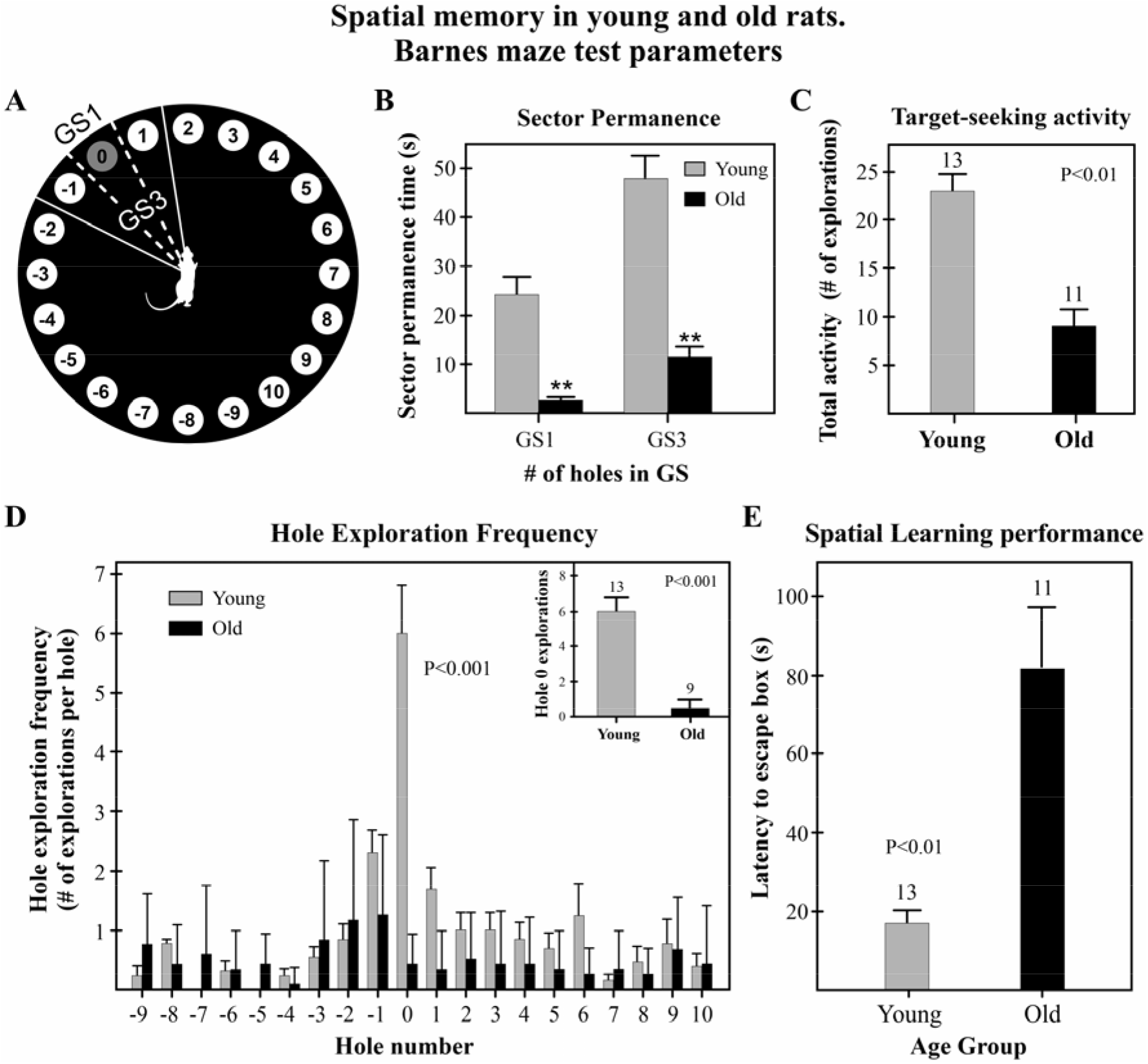
Summary of spatial memory parameters in young and old female rats-. **Panel A-** Barnes maze platform showing the outlines of goal sector 1 (GS1) and goal sector 3 (GS3). **Panel B-** Impact of aging on GS1 and GS3 permanence. Sector permanence is expressed as the time spent in the corresponding sector during the testing time (2 min). **Panel C-** Target (escape box)-seeking activity in young and old animals. **Panel D-** Hole exploration frequency in young and old rats. **Inset-** Number of explorations of hole 0 (escape hole) in young and old rats during the testing time. **Panel E-** Effect of aging on learning and spatial memory retention in rats. Columns represent escape hole latency (seconds it takes for rats to find the escape hole when the box is in place) at the end of the training.

### Chronological age versus DNAm age

While chronological age measures the physical time elapsed since rats were born, epigenetic age reflects biological time, that is the biological dynamics of an organism. A regression analysis of hippocampal DNAm age versus chronological age, showed a very high R value (R=0.99) using testing data **(Fig. 6. panel A)**. The slope was < 1 and regression line intersected the Y axis at nearly 3 months. The same data displayed in a bar chart revealed that in young rats epigenetic time moves faster than physical time (the animals were epigenetically older than indicated by chronological age) but at old ages epigenetic time moves slower that physical time so that while the chronological age of the old group was 26.6 months, their epigenetic ages was only 20.1 months **(Fig . 8 panel B)**. Regression of hippocampal versus blood DNAm age in combined training data from young and old rats revealed a highly significant regression **(Suppl. Fig. 3).**

**Figure 6.**
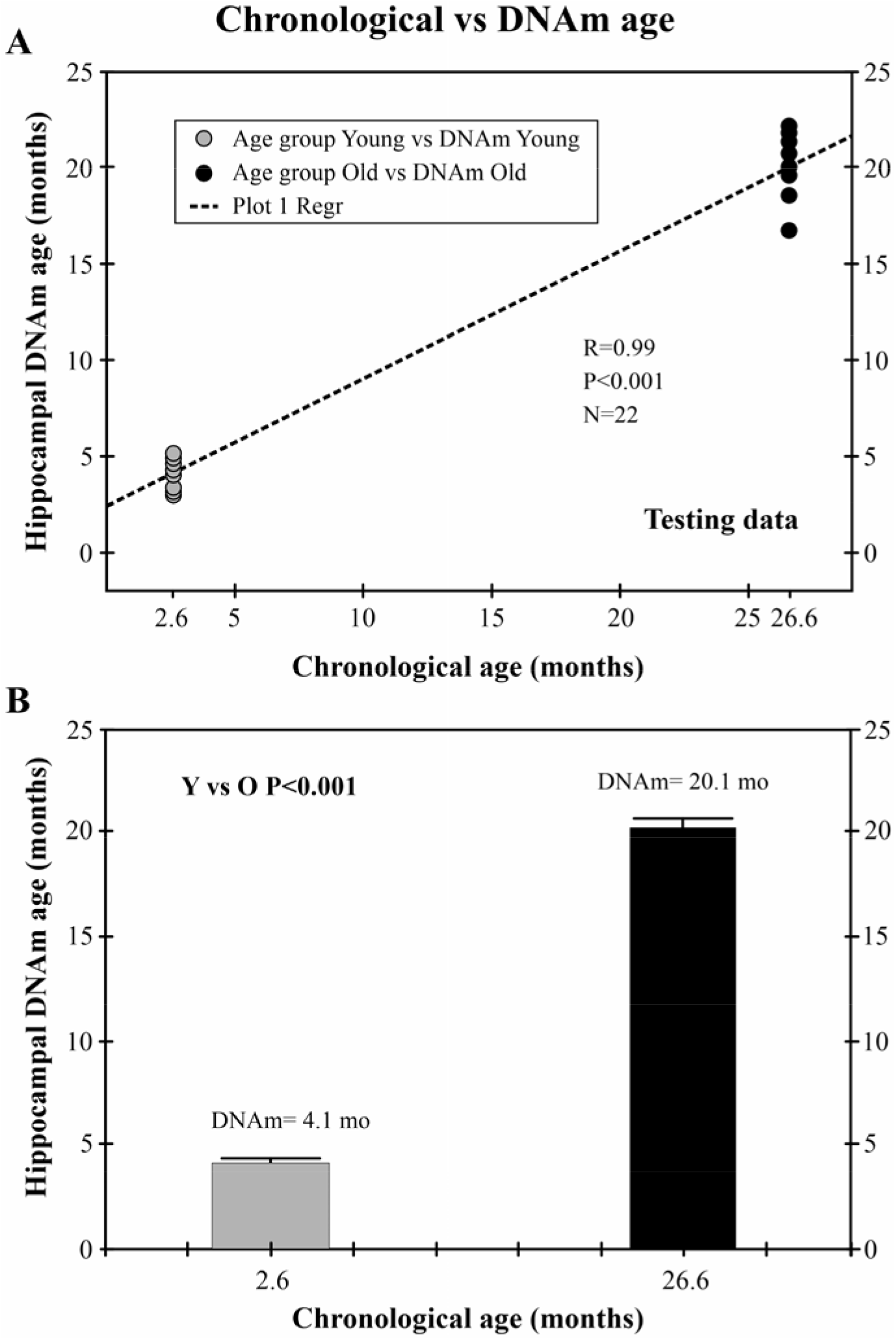
Correlation of hippocampal DNAm age verus chronological age in female rats-. **Panel A-** Regression plot of hippocampal DNA, versus chronological age. Notice the although there is a high correlation between the two types of age, the slope is not =1, that is, they are not identical **Panel B-** The same plot is drawn as a bar chart. Notice that at young ages the epigenetic clock ticks faster than the physical clock. In old animals, the ticking of the DNAm clock is slower than that of physical time.

### Correlation of Barnes maze test indices with hippocampal DNAm age

A regression analysis between Barnes maze test data and hippocampal DNAm age was performed. In all cases, testing data were used for DNAm age. When data were taken from young and old rats together, a significant regression was found for Exploration frequency of hole 0 (R= 0.77). For GS1 and GS3 permanence time, the regression was significant in both cases with R=0.72 and 0.79, respectively **(Fig. 7 panels A, B and C).** When the regressions were done using data from either young or old rats alone, no significant regression was found (data not shown).

**Figure 7.**
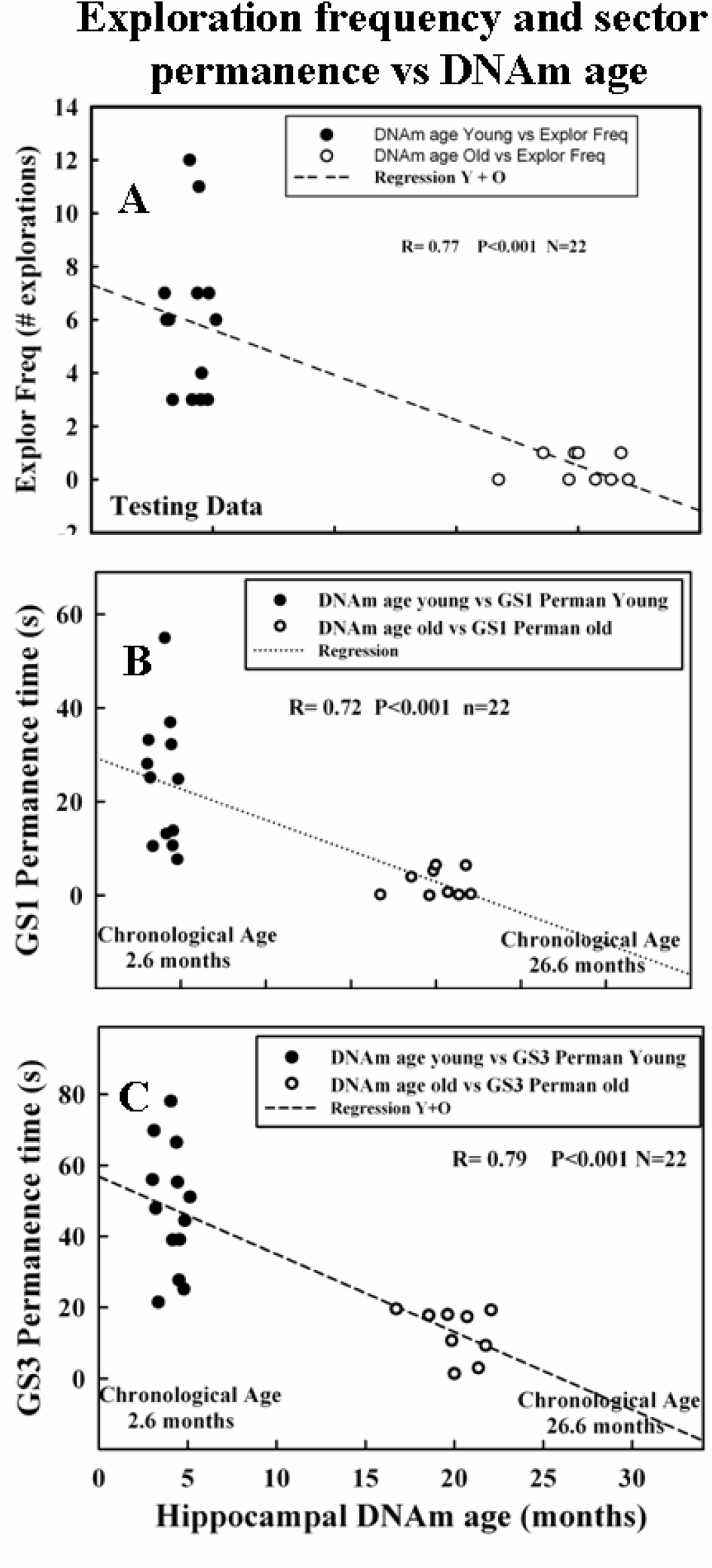
Regression plots correlating spatial memory variables measured by the Barnes maze test versus hippocampal DNAm age. **Panel A-** Regression of hole exploration frequency versus hippocampal DNAm age in young and old rats. The regression is highly significant. **Panel B-** Time spent by rats in goal sector 1 plotted versus hippocampal DNAm age. In this case, the regression is also highly significant. **Panel C-** Same plot for GS3 permanence. The regression also is highly significant here.

**Figure 8.**
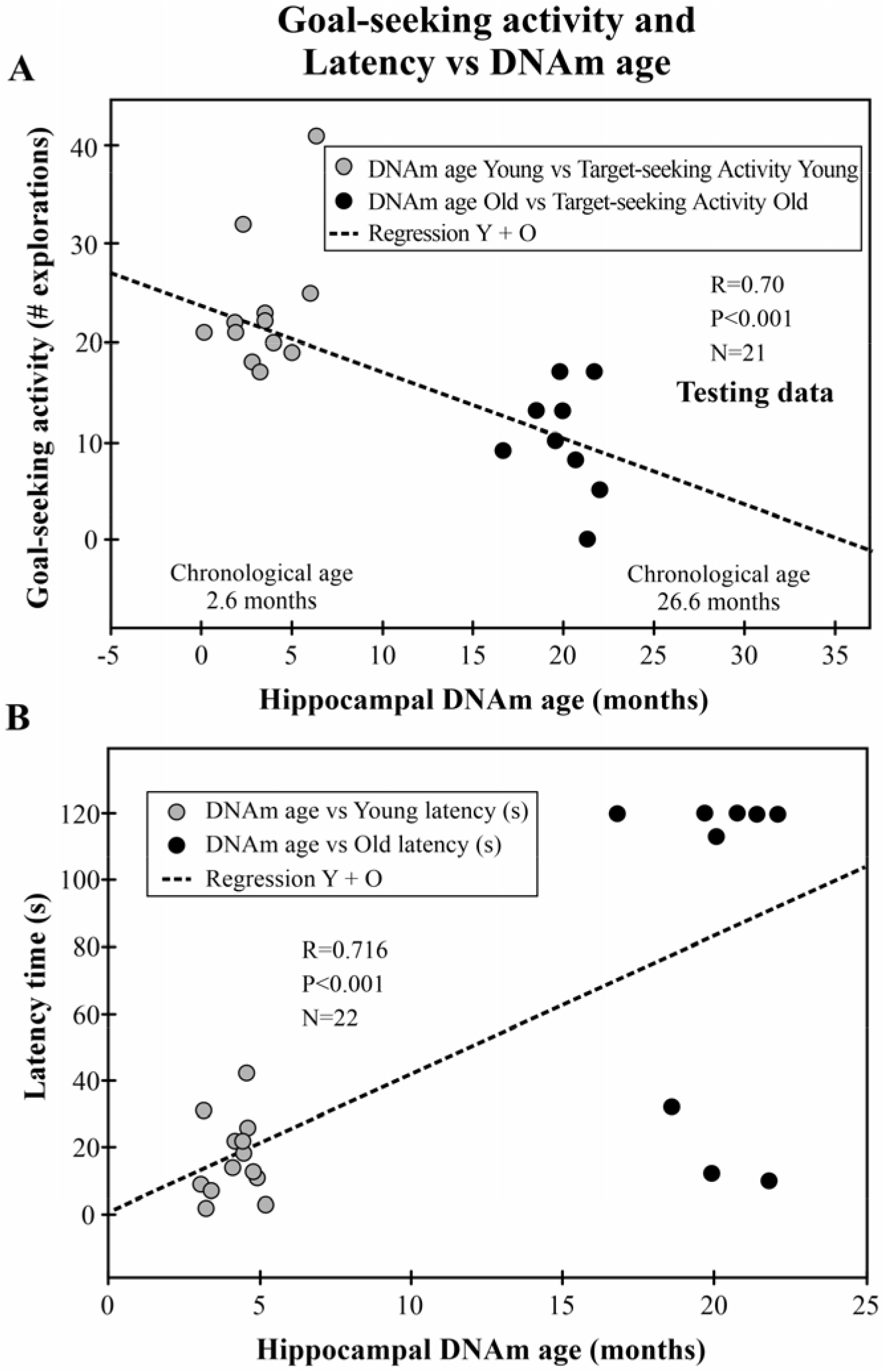
Regression plots correlating cognitive indices versus hippocampal DNAm age. **Panel A-** All-hole exploratory activity versus hippocampal DNAm age. The regression coefficient is highly significant. **Panel B-** Latency time versus hippocampal DNAm age. Latency is an index of learning ability and spatial memory retention. The regression of the plot is highly significant. In all panels, asterisks over bars indicate a highly significant difference from corresponding young counterparts. Numbers over bars indicate the N value for the corresponding group.

Target-Seeking activity showed a significant regression for hippocampal DNAm age with R=0.70 **(Fig. 10, panel B)**. Latency time also showed a significant regression with hippocampal DNAm age with R= 0.72. **(Fig. 10, panel A)**. As before, when the regressions were done using data from either young or old rats, no significance was detected (data not shown).

## DISCUSSION

DNA methylation is one of the best understood modifications of chromatin, with crucial roles in gene expression and imprinting, defense against viral sequences, inhibition of recombination, as well as assembly of heterochromatin **(24).** Three active DNA methyltransferase (DNMT) activities (DNMT1, DNMT3a, and DNMT3b) have been identified in humans and mice **(25–26)**. DNMT1 functions primarily as maintenance DNMT, responsible for copying the parental-strand DNA methylation pattern onto the daughter strand after each round of DNA replication. DNMT3a and DNMT3b function as de novo DNMTs, although they can also maintain methylation patterns. DNA methylation is associated with the induction of a closed chromatin state by recruitment of protein complexes that bear repressive chromatin modifying activities, including MBD (methyl-CpG-binding domain) proteins and histone deacetylase (HDAC)-containing complexes **(27–28)**. Cytosine methylation at euchromatic promoters participates in switching off the corresponding genes. At heterochromatic domains, it participates in the assembly of the highly compacted chromatin characteristic of these domains.

### DNA demethylation in the old rat hippocampus

In the brain, DNA methylation regulates neural activities and memory formation via the control of gene expression in neurons **(29).** Early studies showed that a global loss of DNA methylation occurs in the rat brain during aging **(24).** In line with these finding, our results reveal that in the rat hippocampus aging is associated with predominant DNA demethylation (73% versus 15% hypermethylation). Results in rat brain are consistent with the more general observation that, almost invariably, mammalian aging is more commonly associated with CpG hypomethylation, particularly at repetitive DNA sequences **(30–33)**. This is likely to be at least partly responsible for the loss of heterochromatin during aging. In the hippocampus, aging does not appear to have a significant impact on the methylation levels of CpG islands as only 12% of the differentially methylated CpGs was localized in these DNA regions.

### Increase of hippocampal DNAm in genes of the Zic and Gli families in old rats

In the old rats the results of our GO analysis identified a number of genes that are hypermethylated during aging in the rat hippocampus. In functional terms, the more relevant of the set are the hypermethylated genes of the Zic and Gli families, Zic1, Zic4 and Gli3.

The Zinc finger of the cerebellum genes Zic 1 and Zic4, and the Gli Family Zinc Finger 3 Gli3 play specific roles in development **(34–35).** Zic1, Zic4, and Gli3 gene expression has been found downregulated in the aged hippocampus of Sprague Dawley rats **(36).** It has been reported that the proteins of the Zic and Gli families interact and regulate each other’s transcription through their respective zinc finger domains. Moreover, Zic 1 contributes to the translocation of Gli3 from the cytoplasm to the nucleus **(37)**.

Gli proteins are part of the Hedgehog Signalling pathway, they act as transcription factor activators (or repressors, in absence of the Sonic Hedgehog protein) on the target genes in the last step of the pathway **(38).** Several studies carried out in multiple species concluded that Zic absence can repress Sonic hedgehog (Shh)-signaling, probably by the interaction between Zic and Gli proteins **(39).** Shh signaling was suggested to be involved in age-related neurological disorders since it mediates the formation and plasticity of neuronal circuits **(40).** Thus, our results suggest that the increase in the CpG methylation of these genes in the hippocampus of old rats may have a negative impact on the cognitive performance of the animals, possibly through the effects of the Shh signalling pathway on hippocampal-mediated memory.

### Identification of genes near or at hippocampal GpGs whose methylation levels correlate inversely with GS3 permanence time in old rats

As mentioned above, experience has shown that from the different spatial memory indices the Barnes maze test generates, GS3 permanence time is the one that most reliably quantitates spatial memory performance. For instance, after treatment of 27 months old female rats with naïve or insulin-like growth factor-1 (IGF-1) transgenic human umbilical cord cells, the only Barnes maze indices that detected an improvement in spatial memory in the treated animals were permanence in GS1 and GS3 sectors which increased significantly whereas there were no changes in intact counterparts **(41)**.

Our finding that in old rats of the same chronological age (26.6 mo.) there is a set of 14 hypermethylated CpGs near or at specific genes whose methylation levels are significantly and inversely associated with spatial memory function (as indicated by GS3 permanence time) suggests, although does not prove, that the methylation levels of those CpGs are causally involved in the observed age-related impairment of spatial memory function. The hypothesis is strengthened by the fact that a number of those 14 CpGs that correlate significantly with GS3 permanence time are proximal to genes of the Zic and Gli families. Since the Zic and Gli genes are involved in nervous system development, it is conceivable that during the whole lifetime of rats, the methylation levels of those CpGs increase continuously, decreasing the expression of hippocampal genes important for spatial memory performance. Our results are a descriptive observation in untreated rats.

If an experimental manipulation that slows down aging (i.e., a life extending treatment) were associated with a decrease in methylation levels of those 14 hippocampal CpGs (when compared with untreated controls of the same age), the idea that those CpGs play a causal role in spatial memory performance would be further favored. The observation that the highest R^2^ value, 0.85, observed in this analysis corresponds to CpG 07225115, which is located near the Zic1 gene is in line with the above hypothesis **(Fig. 6 panel J)**.

### Hippocampal DNAm age and spatial memory performance in rats

Considering again the genesis of the epigenetic clock, it is of interest to mention that in 2013 Horvath’s algorithm was successfully tested using approximately 8,000 human DNA methylation data sets from over 30 different tissue types **(11).** For a given chronological age, it was found that in DNA samples taken from whole blood, peripheral blood mononuclear cells, buccal epithelium, colon, adipose, liver, lung, saliva, and uterine cervix, Horvath’s algorithm read essentially the same epigenetic age, the only exceptions being some brain regions and very few other organs **(11, 42, 43).** In healthy humans, the cerebellum is epigenetically younger than the rest of the organism **(44).** Multi-tissue full lifespan epigenetic clocks have been set up for mice **(15, 45)** and more recently, for Fischer344 male rats **(46).**

In a recent study we generated multiple epigenetic clocks for Sprague Dawley (S-D) male and female rats including the design of a pan tissue epigenetic clock for this species **(22)**. Using this new tool, the rate of epigenetic aging of peripheral tissues including liver, ovary, skin, adipose tissue, and blood showed comparable features to those reported in humans and mice **(11, 15, 45)**. The rate of epigenetic aging in a number of brain regions, including the neocortex, cerebellum, hippocampus, substantia nigra, hypothalamus and anterior pituitary were also assessed in S-D females, with the rat pan-tissue clock. Among these six regions, the hypothalamus and pituitary gland showed a marked departure from the average rate of epigenetic aging of the rest of the tissues. In both pituitary gland and hypothalamus, the rate of epigenetic aging was substantially slower than the average rate for other brain regions and peripheral tissues **(22)**. It is of interest that, using training data, blood DNAm age showed a highly significant regression with hippocampal DNAm age **(suppl. Fig. 1).** This is in line with a previous study showing a significant correlation between human brain and blood DNAm modules, which led the authors to conclude that blood is a promising surrogate for brain tissue when studying the effects of age on DNA methylation profiles **(44).**

Our present results reveal that the functional performance of several spatial memory-related indices show significant regression levels when plotted versus hippocampal DNAm age only when data from young and old rats are taken together. In no case regressions of single age groups were significant.

Since the hypermethylation levels of a group of 14 hippocampal CpGs in old rats correlated significantly with spatial memory performance, and epigenetic age is thought to reflect biological rather than chronological age, we expected that regression of different Barnes maze indices versus hippocampal DNAm age in old rats would reveal a significant association. The fact that no regression was significant suggests that hypermethylation of certain hippocampal CpGs may play a more relevant role in memory function than DNAm age.

The observation that hippocampal epigenetic time moves faster than physical time at younger ages but the rate of epigenetic time becomes slower than physical time in old rats is consistent with the fact that developmental changes are faster in young than old animals. The observation is in line with the DNAm age data previously reported in different brain regions of rats of various ages **(22)**.

### CONCLUDING REMARKS

The recent development of the Illumina Infinium HorvathMammalMethylChip40, a mammalian methylation array that possesses a high fidelity in humans, rats, and mice **(17),** has made it possible to profit from a number of advantages that offers the rat as an animal model. For instance its size which is 10 times larger than mouse size is a feature that allows to extract sufficient tissue DNA from small brain regions as the hippocampus. Thus, in rats, we were able to quantitatively assess epigenetic age and other DNAm-related features during aging and rejuvenation **(22)**. In this context, the present results, focused on hippocampal DNAm, show that hippocampal DNAm age is a reliable index of spatial memory performance in young and old rats. Perhaps more significant are the data that come from the GO enrichment and regression analysis in old rats as they suggest that age-related hypermethylation of certain gene families like Zic and Gli may play a causal role in the decline in spatial memory during aging. This is to our knowledge, the first documented study in experimental animals linking age-related changes in hippocampal methylation levels of particular genes with the decline in spatial memory during aging. Our results offer valuable reference values for future epigenetic studies assessing the effectiveness of life-extending or rejuvenation strategies in this species.

## ACKNOWLEDGEMENTS

The authors thank Dr. Kenneth Raj, Public Health England, Didcot, UK, for enlightening discussions on the epigenetic clock and Dr. Martin Abba, CINIBA, UNLP, Argentina for technical comments on methylation analysis. The authors are indebted to Mr. Mario R. Ramos for design of the figures, to Ms. Yolanda E. Sosa for technical and editorial assistance and to Ms. Araceli Bigres for excellent care of our rat colony.

This study was supported in part by grant PICT18—00907, from the National Agency for the Promotion of Science and Technology, Argentina and from research grant MRCF 10-10-17 from the Medical Research Charitable Foundation and the Society for Experimental Gerontological Research, New Zealand to RGG, grant PICT 2018-2446 to CBH and grant #PICT16-1070 to GRM. SH was supported by the Paul G. Allen Frontiers Group.

## Author Approvals

All authors have seen and approved the manuscript, and that it hasn’t been accepted or published elsewhere.

## SUPPLEMENTAL FIGURES

**Supplemental Figure 1.**
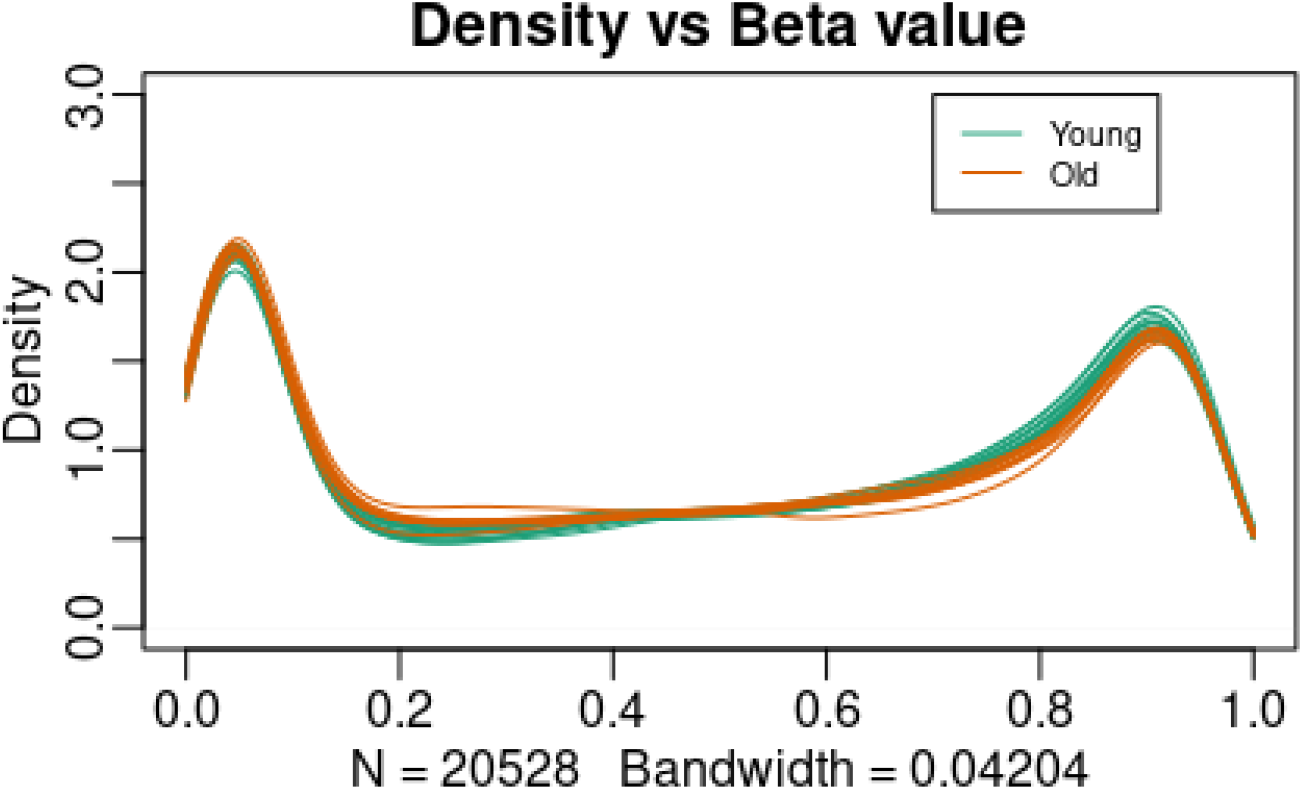
Density distribution of β-values. Density distribution of β-values across all the 22528 CpG sites of the 22 samples. The orange lines represent the distribution of the young animals samples and the green lines represent the distribution of the old animal samples.

**Supplemental Figure 2.**
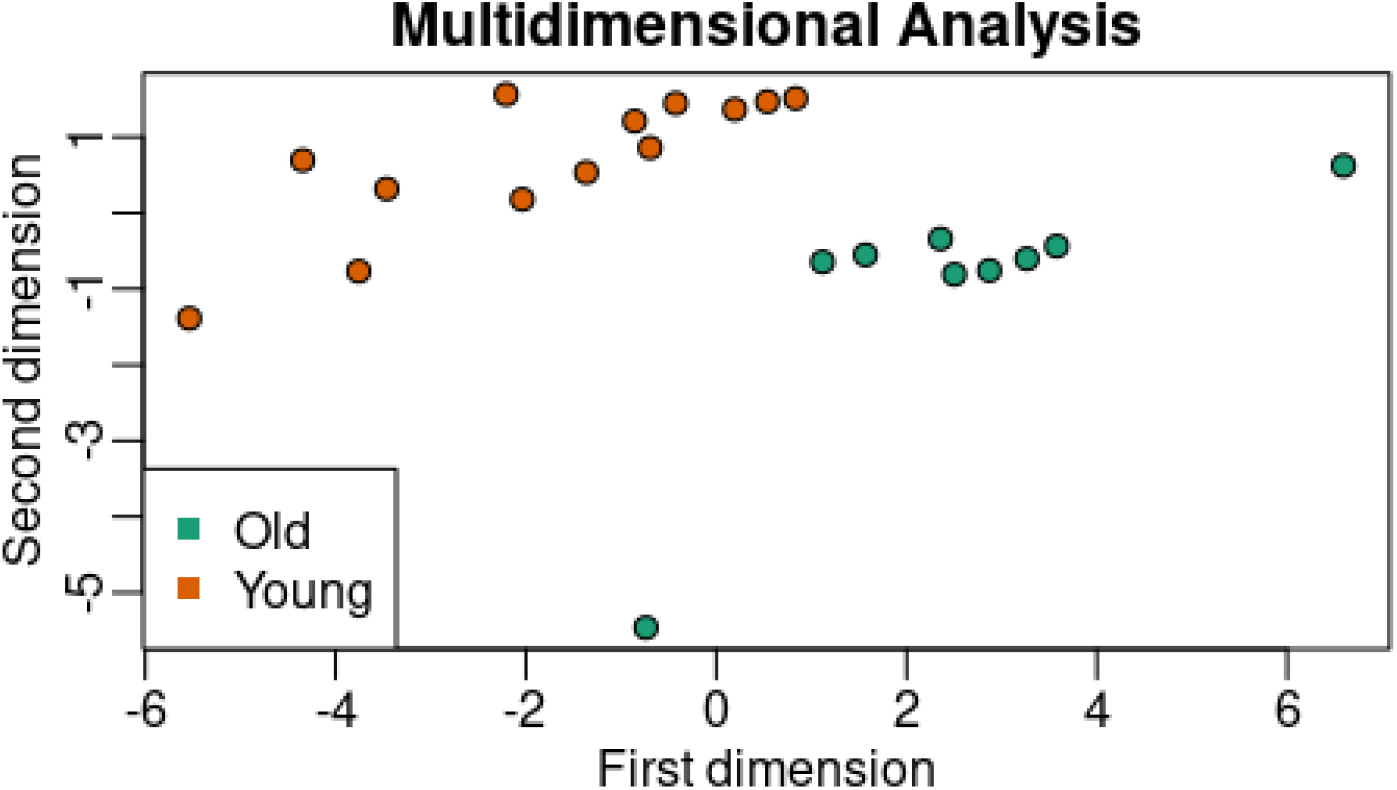
Multidimensional scaling (MDS) plot of DNA methylation profiles in the hippocampus of young and old animals. This MDS plot of 20 samples shows that samples cluster according to age type, as expected.

**Supplemental Figure 3.**
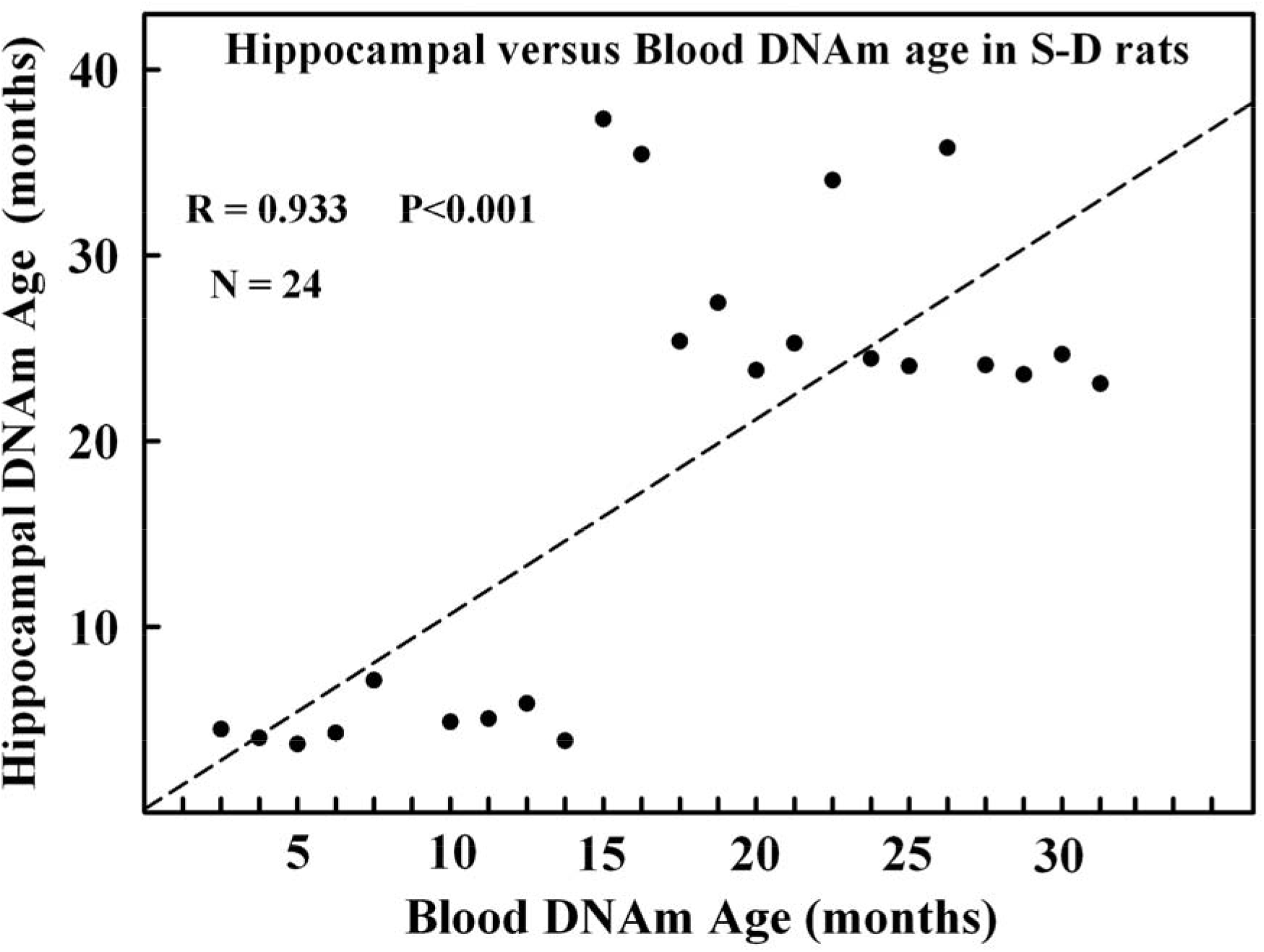
Hippocampal versus blood DNAm age is rats. The regression, using training data, is highly significant which implies that in rats blood DNAm age is a reliable estimate of hippocampal DNAm age.

